# The sum of two halves may be different from the whole. Effects of splitting sequencing samples across lanes

**DOI:** 10.1101/2021.05.10.443429

**Authors:** Eleanor C. Williams, Ruben Chazarra-Gil, Arash Shahsavari, Irina Mohorianu

## Abstract

The advances in high throughput sequencing (HTS) enabled the characterisation of biological processes at an unprecedented level of detail; the majority of hypotheses in molecular biology rely on analyses of HTS data. However, achieving increased robustness and reproducibility of results remains one of the main challenges. Although variability in results may be introduced at various stages, e.g. alignment, summarisation or detection of differences in expression, one source of variability was systematically omitted: the sequencing design which propagates through analyses and may introduce an additional layer of technical variation.

We illustrate qualitative and quantitative differences arising from splitting samples across lanes, on bulk and single-cell sequencing. For bulk mRNAseq data, we focus on differential expression and enrichment analyses; for bulk ChIPseq data, we investigate the effect on peak calling, and peaks’ properties. At single-cell level, we concentrate on identifying cell subpopulations. We rely on markers used for assigning cell identities; both smartSeq and 10x data are presented.

The observed reduction in the number of unique sequenced fragments reduces the level of detail on which the different prediction approaches depend. Further, the sequencing stochasticity adds in a weighting bias corroborated with variable sequencing depths and (yet unexplained) sequencing bias.

## Introduction

The recent developments and improvements in high-throughput sequencing (HTS) technologies facilitated increasingly complex transcriptome/genome-wide analyses [1], enhancing both the qualitative annotation of genomes [2, 3, 4] and their quantitative, functional characterization through differential expression studies [5, 6]. The diversification of methods specialised to a wide range of perspectives on DNA/RNA biology [7] was complemented by studies at single cell level [8]. Advances were observed across all aspects of the sequencing workflow [9], complemented by an increasing amount of resulting data. This created another challenge: producing robust and reproducible results and simultaneously keeping up with the increasing intricacy of experiments [10].

The variability of sequencing output, which propagates through to quantification and other downstream exploration, poses one of the main challenges in bioinformatics analyses, since it implies the disentangling of relevant from irrelevant sources of variation. While the biologically relevant quantities are context dependent [11], an essential distinction exists between variability due to biological processes and variability due to measurement error or inaccuracy [12, 13]. The former is generally specific and well defined in relation to a condition; even when it is perturbed by noise, an underlying pattern of expression may emerge [14]. Technical variability encompasses measurement error [15], sequencing bias [16, 17], and variability due to missing data [18]. For the latter the assessment of technical variation can be hindered by the lack of a ground truth.

Several studies proposed approaches to identify and characterise the sources of variability in HTS experiments, focusing on several aspects of signal distribution, which can affect the accuracy of the downstream analyses and interpretations, and jeopardise the reproducibility of results [11, 19]. These included both the analysis of noise [14, 20] and the downstream components of the analyses such as batch/background effect [21], alignment approaches [22], processing pipelines [5], normalisation methods [23] and differential expression thresholds [24, 25]. To model the intrinsic biological variability, the number of replicates in the context of experimental design was optimised using power calculations [11, 26], designed to provide a robust estimation of differences in expression. These approaches rely on simulations on the number of expressed genes, on mean-dispersion estimates and dropouts after applying frequency and outlier filtering; traditional approaches do not take into account elements of sequencing design including across-lane sample splitting. In general, the impact of library construction and flow cell and lane characteristics on downstream analysis has not been studied in detail.

Sources of technical variability for RNAseq experiments span from the combinatorial numbers of highly variable isoforms to the handling of ambiguous or multi-mapped reads [1]. For ChIP datasets, the ability to address specific biological questions can be significantly impacted by antibody efficiency and specificity [27] as peak distributions are a direct consequence of affinity, over-crosslinking, DNA fragmentation and PCR amplification; for such samples, users are faced with a trade-off between number of usable reads (sensitivity of peak detection) and proportion of false positives derived from multi-mapped reads [28]. Low quality replicates can also generate bottlenecks when used in conjunction with good samples, as true peaks missing from poor quality replicates may be marked as nonreproducible, thus creating false negatives [27]. Single-cell experiments share some of the drawbacks of bulk ones; in addition, the exponential increase in the number of cells profiled per study, coupled with the shallower sequencing depth, redefined some of the known difficulties, such as the characterisation of noise [29, 20].

Here we investigate the effect of across-lane sample splitting, at the sequencing stage, on downstream analyses for bulk and single cell data; the sampling approach is modelled on observed sequencing outputs (bulk mRNAseq data). To infer the effect on other types of sequencing data, we study the differences between ground truth and split-samples (simulating across-lane splitting), with various parameters controlling the number of splits and proportion of reads produced on each lane. We focus on standard analyses i.e. identification of differentially expressed (DE) genes or ChIP peaks for bulk analyses; for single cell analyses, we concentrate on the allocation of cells to clusters (viewed as proxies for cell types), and comment on the observed variability in biological interpretations.

## Methods and Materials

### Materials

The motivation for the subsampling strategy used throughout the manuscript is derived from a *D. melanogaster* mRNAseq dataset (GSE85806) for which 3 samples were sequenced split across 2 lanes (GSM2284703, GSM2284704 (2RA3), GSM2284705, GSM2284706 (2RH2), GSM2284707, GSM2284708 (26RH3)). To highlight the consequences of this choice in sequencing design we compared the resulting expression levels to the corresponding full-samples, GSE55839 (GSM1346985 (2RH2), GSM1346996 (2RA3), GSM1347001 (26RH3)) [30].

To illustrate the split effects and their link to the biological interpretation we use bulk and single-cell mRNA data, and bulk ChIPseq data. For the former (bulk mRNAseq) we used the Yang et al 2019 dataset [31], focusing on the 0h and 12h samples (GSE117896, comprising SRR7624365, SRR7624366 (biological replicates for 0h), SRR7624371 and SRR7624372 (biological replicates for 12h)). The bulk ChIPseq analysis was performed on H3K4me3 and H3K27ac samples, using 0h and 12h samples for each (SRR7624381, SRR7624384, SRR7624389, SRR7624392).

To exemplify the effect on plate-based scRNA-seq platforms (smartSeq) we use the Cuomo et al 2020 dataset [32]. We selected data from 6 donors, on 4 time points. On this input six experimental study cases (Table 1) were designed to illustrate the effect of the different covariates i.e. the donor, the specific time-point and cell types (resulting clusters) identities. To investigate the effects of lane-splitting on 10x Genomics scRNA-seq data, we used an in-vivo dataset of human hematopoietic stem and progenitor cells from spleen, bone marrow, and peripheral blood [33]; the data is available via BioStudies accession S-SUBS4 (donor SAMEA6646089).

**Table 1:** Overview of the study cases built on a scRNA-seq Smart-seq data (Cuomo et al 2020), illustrating the combinations of the different covariates. Study case 4 (same donor, different time-point and same cluster) and case 7 (different donor, different time-point, and same cluster) could not be generated due to the data structure.

|  | Study case | N cells | Donor | N cells (donor) | Time point | N cells (time point) | Cluster | N cells (cluster) |
| --- | --- | --- | --- | --- | --- | --- | --- | --- |
| 1 | Case 1 | 105 | hayt | 105 | day2 | 105 | 0 | 105 |
| 2 | Case 2 | 106 | pahc | 106 | day3 | 106 | 1 | 66 |
|  |  |  |  |  |  |  | 4 | 40 |
| 3 | Case 3 | 168 | melw | 94 | day0 | 168 | 3 | 168 |
|  |  |  | qunz | 74 |  |  |  |  |
| 4 | Case 5 | 168 | hayt | 168 | day1 | 61 | 9 | 61 |
|  |  |  |  |  | day3 | 107 | 1 | 107 |
| 5 | Case 6 | 95 | melw | 47 | day1 | 95 | 5 | 45 |
|  |  |  | vils | 48 |  |  | 6 | 50 |
| 6 | Case 8 | 217 | melw | 95 | day0 | 95 | 3 | 95 |
|  |  |  | naah | 122 | day3 | 122 | 2 | 122 |

### Methods

#### splitting strategy - sequenced samples

For the 3 *D. melanogaster* samples (2RA3, 2RH2, 26RH3) for which whole-lane and split-lane sequencing was available, we followed the standard mRNAseq quantification procedure; the split samples were merged, without any additional pre-processing (merged-samples). Whole, split and merged samples were aligned to the *D. melanogaster* r6.41 genome [34] using STAR 2.7.0a [35], with default parameters. Next, the expression was quantified using featureCounts 2.0.0 [36], and summarised into count matrices. For each BAM, a bigwig was produced using bamCoverage and individual transcript coverage was identified using pyBigWig from deeptools [37]. In addition, for all settings, we determined the number of non-redundant (unique) and redundant (all) reads and evaluated the number of fragments present exclusively in the one setting. We also calculated the ratio between the abundance of a read (its redundancy) in the whole vs split sample, with an expected value equal to the ratio of sequencing depths.

#### splitting strategy - simulated data

The splitting strategy for the simulation study is consistent across all datasets. The splitting is performed per sample. For each dataset, on the ground truth (GT), i.e. the original sample, and on the simulated, split (S) samples, the same pipelines (for alignment, quantification and identification of differentially expressed entries), with identical, default parameters are applied, to ensure an unbiased comparison between results obtained on the GT and S samples, respectively.

Let *k* be the number of splits for a sample, and *n* the number of iterations (each split sample is generated from the original GT sample). The steps for generating an S sample are: [1] subsampling reads without replacement from the GT sample to 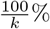 of GT in *n* iterations; [2] subsequently the *k* subsamples are concatenated. For bulk mRNAseq and ChIPseq samples, we assessed *k* = 2 and *k* = 3, *n* = 10. In Discussion, we present simulated samples for *k* = 2, 3, 4, 5 for one GT sample (bulk mRNAseq, 0h rep 1), *n* = 10; we also analysed simulated samples for *k* = 2 with variable split proportions from 55 - 45 to 90 - 10. For the bulk ChIPseq data we assessed *k* = 2 and *k* = 3, *n* = 10. For the smartSeq data and 10x data we generated *n* = 3 S samples for *k* = 2, for each of the study cases (i.e. each subset of samples), respectively, using the seqtk toolkit (https://github.com/lh3/seqtk). We note that due to the stochasticity of sequencing we cannot simulate fragments that are present exclusively in the whole or split samples. This limitation of the simulation study is compensated by the analysis of true split samples (the *D. melanogaster* dataset).

#### bulk mRNAseq, *Yang et al* dataset

The raw samples (no QC-based filters applied) were aligned to the *M. musculus* genome [Ensembl 98.38] [38] using STAR 2.7.0a (paired-end mode) [35]. Next, the expression was quantified using featureCounts 2.0.0 [36]. To assess the stability between S and GT samples, abundance density plots and MA plots were produced. We used noisyR [14] to perform noise analysis on the GT and S samples, and further analysed the PCC distribution for binned abundances. For each dataset, expression levels were normalised using quantile normalisation [39] and DE genes between 0h and 12h were identified using edgeR [40]. The DE call was based on |*log*_2_FC| > 0.5 and adjusted p-value < 0.05 (using Benjamini-Hochberg multiple testing correction). Enrichment analysis on the resulting DE genes was performed using the gprofiler2 package [41] on GO, KEGG, reactome and transcription factor terms, with the background set as the set of expressed genes with abundance > 0 in at least one sample.

#### bulk ChIPseq

The GT and S samples were aligned to the *M. musculus* genome using STAR 2.7.0a (single-end mode using default parameters) [35]. Narrow peaks were called using macs2 2.2.7.1 [42]. Peaks were matched between samples if the midpoint of the peak in sample 1 is within the boundaries of the peak in sample 2 and vice versa. Across sets of samples, amplitudes were normalised using quantile normalisation [39] and differentially methylated peaks between 0h and 12h were identified using edgeR [40]. Peaks were called DE if |*log*_2_FC| > 0.5 and adjusted p-value < 0.05 (using Benjamini-Hochberg multiple testing correction).

#### sc smartSeq

Six experimental study cases were designed based on the Cuomo dataset [32], considering the donor, time-point and cluster cell identities (Table 1). The corresponding samples were standardised in read length using Trim Galore (0.4.1) by: trimming 10bp from the 5’ end (to reduce the effect of sequencing bias) and 40 bp from the 3’ end (to address high adapter sequence content); the length of the resulting reads was 75 nts. The GT and S samples were aligned to the *H. sapiens* genome (GRCh38.p13) [38] using STAR (2.7.0a) [35] in paired-end mode. The gene counts were summarised in a matrix using featureCounts 2.0.0 [36]. We applied fastQC [43] to obtain read quality metrics and multiQC [44] to aggregate QC results from the reads, alignment and quantification. Seurat objects [45] were created considering features expressed in > 3 cells, and cells with > 50 features. The analysis pipeline comprises: i) normalization of expression levels (SCTransform [46]), ii) computation of PCA and UMAP embeddings (RunPCA, RunUMAP), iii) Neighborhood graph computation (FindNeighbors), iv) Clustering (FindClusters), v) Differential Expression (FindAllMarkers). We considered as DE features those with log_2_ FC > 0.5 (i.e. the positive markers) and an adjusted p-value < 0.05 (using Bonferroni multiple testing correction). To assess the similarity of the partitioning in the GT and S samples, we calculated Jaccard similarity indices (JSI, [47]) on the cluster-specific sets of DE features (Fig 2D); restricting the JSI to the smaller set of markers.

#### sc 10x Genomics

The GT and S fastq files were aligned to the 10x *H. sapiens* GRCh38 v3 reference transcriptome; the protein-coding genes were quantified using 10x Cellranger v3.1.0 [48]. The processing and analysis was performed individually for each donor, for the GT and S samples: after inspection of distributions of UMIs, number of detected genes, proportion of UMIs from mitochondrial genes (MT) and ribosomal protein-coding genes (RP), only cells with > 1, 000 unique genes, < 10% MT and > 20% RP were retained for downstream analysis; the MT and RP genes were subsequently discarded from the count matrix. Following normalization using SCTransform [46], the 3,000 most abundant genes, accounting for 60% − 85% of UMIs in cells across the data, were identified and used for the calculation of PCA. A UMAP dimensionality reduction was calculated using the 30 first PCs (after inspection of an elbow plot of PC variance); the UMAP was subsequently used to assess the extent of potential batch effects originating from the tissue origin of cells, raw and normalized sequencing depths, MT% and RP%.

A 20-nearest neighbour graph was computed on the first 30 PCs of the data; the cells were clustered using SLM community detection [49] on the NN graph. To assess the clustering similarity, element centric clustering comparison [50] was employed on the set of common barcodes between GT and S using the ClustAssess R package [51]. Cluster markers were identified using the ROC test in Seurat v3.1.4 [45]; only genes with |FC| > 2 were considered. The top 25 markers per cluster, ranked by discriminative power, were subsequently used to calculate the per-cell JSI between cluster markers; the JSI when at least one of the sets is empty was set, by default, to 0.

All analyses were performed in R (3.6.3). The code for generating these results is available on github https://github.com/Core-Bioinformatics/split-manuscript. All tests were performed on a Linux server (16 cores, 755G RAM).

## Results

### Across-lane split leads to differences in bulk mRNAseq data

To determine (and justify) the appropriate parameters for the simulation study, we analysed the properties of 3 *D. melanogaster* bulk mRNAseq samples for which both the whole-sample per lane [30] and a quantification-based 50/50 split output were available. We note that the sequencing depths for the halves-samples (2RH2: 36M/26M (58%/42%), 2RA3: 21M/31M (41%/59%), 26RH3: 33M/36M (47%/53%)) diverged from the expected 50/50. To assess the consistency in properties for split-samples vs whole-samples, we first evaluated the nucleotide composition; we see no significant (BH-adjusted p-value < 0.05) differences when comparing, per base, nucleotide distributions or GC content (assessed using *χ*^2^ tests on A/C/G/T frequencies per position, Supplementary table 1). Next, we looked into the number and abundance of non-redundant (unique) fragments; we illustrate the distributions of abundances for the specific fragments, across the 3 available samples.

Although we see an increase in sequencing depth for the merged-samples vs the whole-samples, we do not consistently observe a similar proportional increase in the number of unique reads; for 2RA3 the ratio of unique reads in the merged-samples compared to the whole is ∼ 1.16 (compared to ∼ 1.36 ratio on sequencing depths), we observe a similar proportion for 2RH2 (∼ 1.17 compared to ∼ 1.18) and for 26RH3 (∼ 1.20 compared to ∼ 1.23). Next, we assessed the ratio between the abundances of individual reads in the whole-samples vs the merged-samples, the analysis was performed on non-zero counts for both compared samples; the ratio is expected to mirror the proportions of sequencing depths. In Supplementary fig. 1 A-C, we show the distribution of log_2_ ratios of abundances between merged- and whole-samples, binned on abundances; the guide line indicates the expected sequencing-depth-based ratio. The ratio distributions are located below the guide line for low-medium abundances (1-6 on the log_2_ scale); the ratio distributions increase to the expected level for medium range abundances, and rise above the guide line for high abundances, underlining an “over-amplification” of signal and a sensitivity of signal quantification to the lane splitting strategy. Most specific fragments (to either the concatenated samples, or the whole ones) are low abundance; however, we note a few high abundance fragments which may be affected by sequencing bias (the distributions of abundances are summarised in Supp Fig 1, D-F).

The change in ratio between split- and whole-samples is also reinforced when we consider the differences between the average abundances, calculated using all incident reads, on sliding windows (100nts) for the whole- and merged-samples (Supp Fig 1 G-I). The ratios are scaled per overall window abundance. Positive differences (red) correspond to higher amplitudes in the whole-sample; negative differences (blue) correspond to higher amplitudes in the merged-samples). For each sample, we show both the distribution for all differences (top subplot) and for differences above 0.2 (bottom subplot); the number of windows in each abundance bin are shown above the corresponding boxplots. We see wider variation between whole and merged-samples at low-medium abundances (0-7 on the log_2_ scale) with larger differences when the merged-sample abundance is higher than one in the whole sample. This trend suggests a systematic, consistent stochastic over-amplification in the merged-samples; conversely, for abundances higher in the whole sample, the differences in the merged-samples are more subtle. For medium-high abundances (> 7, log_2_ scale), we see a higher number of small differences for windows with abundances higher in the merged-sample; the range of differences is wider for for windows with higher abundance in the whole-samples, but their count is lower. The pre-alignment analysis, at read level (Supp Fig. 1 A-C) hinted to these more subtle downstream effects, underlying the importance of studying the impact of splitting across lanes on real data and simulated case studies.

To illustrate some differences observed on individual transcripts, we present three examples (from sample 2RH2) of expression profiles, quantified using the all reads from the whole, the split and the merged samples, for transcripts at high, medium and low abundances; these examples were selected based on the differences in signal distributions. The two split samples behave as technical replicates, as expected; however for the merged sample, we observe significant variation in distribution of signal compared to the whole sample (panel L), or in the localisation of expression features such as peaks (panels J, K)). These differences may have knock-on effects on the downstream analyses and interpretation results (e.g. for the quantification of noise [14]).

These remarks on sequenced output generate the hypothesis that conclusions drawn from standard comparisons between samples may be altered by variations in splitting strategies; this effect is also entirely technical and stems from the stochasticity of sequencing. While the simulation approach proposed in this study cannot capture fragments found exclusively in split samples, the subsampling/concatenating process mirrors the under-representation of low-medium fragments and over-amplification of high abundance fragments, as well as loss of read diversity.

### Consequences of across-lane splitting on bulk data

To systematically investigate the consequences of the observed differences resulting from splitting samples across lanes, we simulated the splitting of sequencing samples on several datasets and assesses the downstream effects.

#### bulk mRNAseq

For the bulk mRNAseq study case [31], we focus on the effect of across-lane sample splitting on expression quantification, DE calling and enrichment analysis (summarised in Fig 1A-F). First, we assess the differences in quantification corresponding to the same time-point/biological replicate, 0h (rep 1), on GT vs *k* = 2 S sample S_2_ (1A) and *k* = 2 vs *k* = 3 samples S_2_ and S_3_ (1B). We observe standard MA funneling shapes, with a variation in excess of 1.5FC for low abundances for both comparisons. In addition, for low to medium abundances, the *S*_2_−*S*_3_ comparison yields a wider distribution of log_2_ FCs underlining the technical variation (noise) that is introduced (Fig 1B). Also noted is a slight shift of FCs (Fig 1A) driven by the stochastic redistribution of reads i.e. lack of signal for some low abundance genes, and over-expression of medium-high abundance genes. Similar conclusions are presented in Supplementary Fig 2A and 2B illustrating GT vs *k* = 3 samples S1_3_ and two simulations of *k* = 2 S1_2_ and S2_3_, respectively.

**Figure 1:**
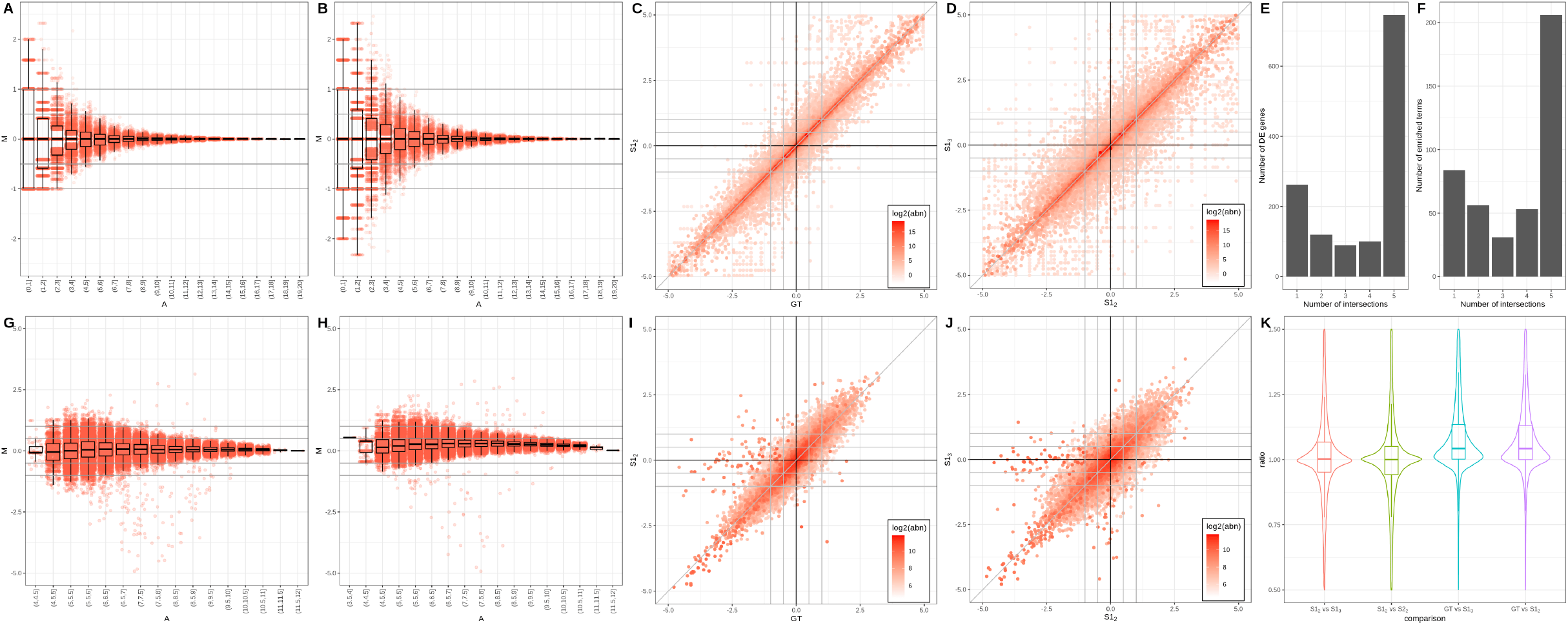
Comparison of bulk mRNAseq analysis results for GT and S samples. A-B. Scatter (MA) and box plot summary on GT sample (0h, rep1) and a corresponding *k* = 2 S sample S1_2_ (subplot A) and *k* = 2 and *k* = 3 S samples S1_2_ and S1_3_ (subplot B). C-D. Scatter (cross) plots comparing the DE amplitude (log_2_ FC) calculated on 0h vs 12 h samples for the GT vs *k* = 2 S samples S1_2_ (subplot C), and *k* = 2 and *k* = 3 S samples S1_2_ and S1_3_ (subplot D); the colour gradient is log_2_-proportional to the average abundance across all 8 corresponding samples. E) Upset plot showing intersections between sets of DE genes (0h vs 12h) in GT samples (GT), 2 sets of *k* = 2 S samples (S1_2_ and S2_2_) and 2 sets of *k* = 3 S samples (S1_3_ and S2_3_). F) Upset plot showing intersections between sets of significant terms (adj p-val *≤* 0.05, BH correction) predicted using gprofiler2 on the corresponding DE genes (see E). **Comparison of H3K4me3 ChIPseq analysis results for GT and S samples**. G-H. Scatter (MA) and box plots showing log_2_ abundance against log_2_ FC within G) GT sample (0h) and a *k* = 2 S sample S1_2_ and H) *k* = 2 and *k* = 3 S samples S1_2_ and S1_3_ for the same sample (0h) I-J. Scatter (cross) plot showing log_2_FC when comparing I) 0h and 12h for GT samples and *k* = 2 S samples S1_2_, and J) 0h and 12h for *k* = 2 and *k* = 3 S samples S1_2_ and S1_3_, coloured by the average amplitude across all 4 corresponding samples. K) Violin and box plots showing distribution of ratio of peak lengths between 2 samples 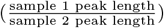, comparing ground truth (GT), *k* = 2 S sample (S1_2_ and S2_2_) and *k* = 3 S samples (S1_3_).

Next, we focus on the DE call between the 0h vs 12h replicates, determined using standard pipelines and parameters. We illustrate the properties of the DE sets called on GT vs *k* = 2 (Fig 1C) and *k* = 2 vs *k* = 3 (Fig 1D), respectively; overall the results converge i.e. all genes are located in the proximity of the diagonal, however the variation is larger for the *k* = 2 vs *k* = 3 simulation S1_2_ vs S1_3_ (1D). Specifically, we see 10,810 genes (31.2% of expressed genes) with > 0.5 log_2_FC absolute difference for S1_2_ vs S1_3_, compared to 7737 (22.3%) between GT and S1_2_. The colour gradient is proportional to the average abundance of genes and underlines that medium abundance genes are mostly affected by the variability in fold-change amplitude and identity. We also assessed the stability of DE signal, for the GT and two sets of simulations for *k* = 2 and *k* = 3, summarised as an upset plot (the *x/*5 intersections are represented, with *x* ∈ {5, 4, 3, 2, 1}). While the majority of genes are detected across all comparisons (746 out of 920 for DE in GT and 1318 called DE in any GT or S sample), we notice comparison-specific genes (up to 81 out of 1012 called DE for S2_3_). In particular, this analysis revealed only few “false negatives” (4 genes DE only in GT samples) in contrast to a larger number of “false positives” (between 48 and 81 from each S sample). 1 gene (out of 1318 called DE in any GT or S sample) was identified across all S samples but not in the GT. To link back the variability of signal to the biological interpretation of the results, we compare the sets of enriched terms, corresponding to the various DE predictions (Fig 1F). Similarly to the DE summary (Fig 1E), we observe a 68.3% consistency in results i.e. significant terms shared between all comparisons (out of 101 identified in total for GT, 141 identified for at least one comparison). However, “false positive” entries are present (24 in total, between 2 and 11 from each S sample) identified only for one set of S samples. These corroborated results underline a mixture of consistent patterns (on S samples) and random variation, difficult to predict or mitigate. This suggests lane-splitting introduces non biologically-robust results.

#### bulk ChIPseq

The effects of across-lane sample splitting are also observed on ChIPseq data; for this case-study, we will focus on the changes in peak amplitude and length; H3K4me3 (Fig 1G-K and Supplementary Fig 2E-J) and H3K27ac (Supplementary Fig 2K-Q) data will be exemplified [31]. The variability on peak calling for H3K4me3 data is assessed using MA (Fig 1G,H) and cross plots (Fig 1I,J). Similarly as for the mRNAseq data, we observe a funneling behaviour when GT and S samples are compared (Fig 1G and H), a wider variation in FC for S1_2_ vs S1_3_ (Fig 1H), and a over-amplification of peak abundances in the simulated samples (Fig 1G); additional supporting results are presented in Supplementary Fig 2E,F. These fluctuations propagate on DE calls (performed using standard pipelines, on the 0h and 12h replicates), and are summarised in Fig 1I and 1J. Despite the overall convergence, for the H3K4me3 data we notice a wider variability than for the mRNAseq data, with clear “false negative” peaks in the simulated data i.e. medium abundance peaks identified as DE in the GT comparison, with a log_2_ FC > 2, and not detected as DE in the S samples (borderline vanishing peaks); this behaviour observed also for H3K27ac data (Supplementary Fig 2M,N).

To further investigate the reduced robustness for the DE call, we focused on other peak properties such as peak length (defined as stop - start coordinates from the macs2 narrowPeak output); in Fig 1K we illustrate the distribution of ratios of peak lengths when GT and S samples are compared. We notice a global, systematic shift towards shorter peaks in simulated samples, the effect is more pronounced for *k* = 3, and a stability of behaviour across simulations (blue and purple distributions). The decrease in peak length can only be observed for common peaks. The reduction in number of called peaks is illustrated in Supplementary Fig 2I,J. For the 0h sample (SRR7624381), we see 10846 fewer peaks called for S1_2_ samples (mean across 10 iterations) compared to GT and 14885 fewer for S1_3_ samples compared to GT, yielding percentage decreases averaging 12.7% and 18.4% respectively.

Simulated across-lane sampling splitting has multiple effects on the peaks called in ChIPseq data, from the number of peaks to their amplitude and length; this technical, stochastic variation may have an impact on the interpretation of results.

### Effects of across-lane splitting on single-cell data

#### sc smartSeq

We exemplify the effect of across-lane sample splitting, on single-cell data, first on a smartSeq2 dataset. The diversity of conditions (donors and time-points, illustrated in Fig 2A) of the Cuomo et al 2020 dataset [32] enables us to incrementally examine the consequences across 6 study cases (Table 1), covering a wide range of experimental situations.

**Figure 2:**
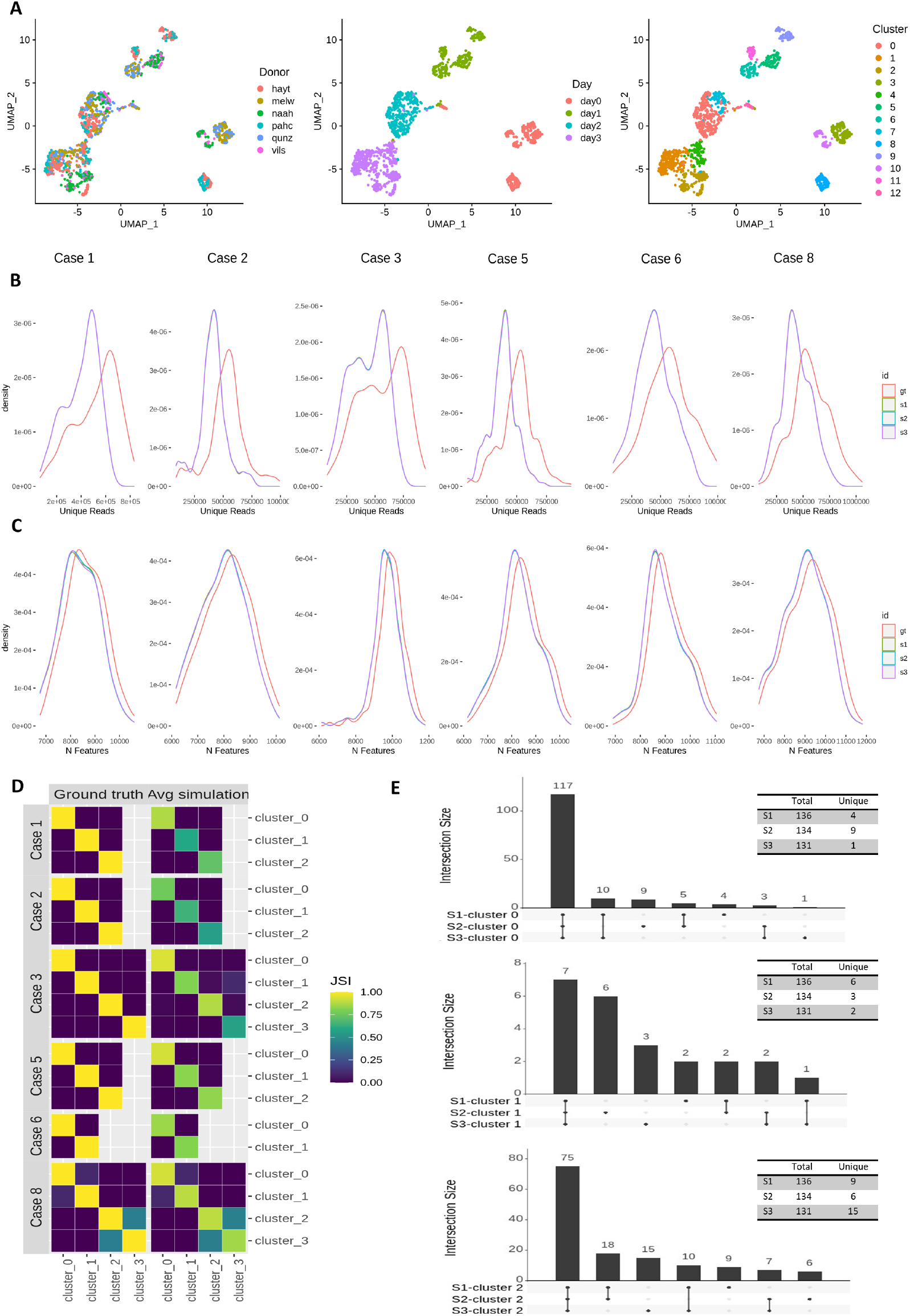
Sample splitting across sequencing lanes introduces variability that is propagated to downstream analysis in Smart-seq2 scRNA-seq data. A) UMAP representation of the Cuomo dataset. Cells are coloured according to their donor (left), time-point (central) and cluster (right). For the different study cases considered we show: B) Number of Unique Reads distribution for the ground truth and simulations. C) Number of features distribution for the GT and S samples. D) Cluster similarity for each study case as evaluated using the JSI calculated on the set of DE features obtained per cluster. The values across the 3 simulations were averaged using the geometric mean. E) Variability of differentially expressed feature set and number of unique DE features across simulations for cluster 0 (upper), 1 (central) and 2 (lower) for Study Case 1.

At the sequencing read level, the sample splitting significantly reduces the number of unique reads (Fig 2B, Supp Table 2A) i.e. the read diversity decreases due to the expansion (over-representation) of duplicated reads. On an invariant total number of reads, this decrease propagates through to the subsequent alignment and quantification steps without influencing the fraction of mapped reads and the count distribution of samples, respectively (Supp Fig 3B,C); nonetheless, this results in a reduction of the number features detected in S samples (Fig 2C). This is a consequence of the loss of low-abundant features, while mid-abundance and high abundance features slightly increase their expression due to the higher read duplication. Significant differences in the number of features (< 0.01 type I error) between simulations and ground truth are detected only for study Case 3; the abundance distributions are not significantly different (in localisation or shape) across the other study cases (Supp Tables 2A,B). Despite this, the reduction in the number of observed features has an impact on downstream analyses such as partitioning into clusters and cell visualization or identification of cell type markers. For the former, the UMAP topology of cells/ clusters is altered (Supplementary Fig 3A); for the latter, in Fig 2D we observe a generalised reduction in the similarity of marker genes for S samples when compared to marker genes determined on the ground truth sample (overall, 0.55-0.94 of the markers are shared between homologous clusters in GT and S samples). In addition, to assess the variability across simulations, we display summary upset plots of marker genes across clusters and simulations (Fig 2E, Supplementary Fig 3E). Although the majority of the DE entries are shared between simulations, unique features are present in most cases; these sum up to 12.3% of total features, on average, across clusters, simulations and study cases.

Overall these results suggest that sample splitting reduces read diversity, which is propagated to downstream analyses, altering the number of features expressed in samples, and potentially altering the biological interpretation of results. Moreover, we highlight the variability across simulations which may introduce an additional degree of irreproducibility across replicates.

#### sc 10x Genomics

The second single-cell case study focuses on a 10x dataset [33]; a similar strategy for generating split samples was applied. Following the alignment and protein-coding gene quantification of the GT and S samples, we observe a high overlap of called cells (i.e. barcodes) resulting from the Cellranger cell calling algorithm (19,453 common barcodes, 166 barcodes unique to ground truth, 13 barcodes unique to simulated). After filtering the low-quality cells, the vast majority of barcodes were still common to both versions of the analysis (17,095 barcodes in common, 155 barcodes unique to GT, 5 barcodes unique to simulated). We observe a greater diversity of UMIs and genes in GT compared to simulated samples (Supplementary Fig 5F-G).

Following SLM clustering [49] on the nearest-neighbor graph, 19 clusters were found on GT and 18 on the simulation of P1 (Fig 3A,B). Element-centric clustering comparison [50] was used to evaluate the per-cell clustering similarity on the set of barcodes common to GT and simulated samples (Fig 3C), revealing high similarity (ECS > 0.6) for 91% of cells (the top of the GT UMAP) and especially the for the island on the upper right (cluster 4 in both clusterings); low similarity (ECS < 0.2) was observed for 9.0% of cells (mainly bottom of the UMAP); no cells had intermediate ECS (0.2 ≤ ECS ≤ 0.6). Notably, an island of cells at the lower right of the GT UMAP, corresponding to cluster 15 in GT, disappeared in the split lane simulation (S samples).

**Figure 3:**
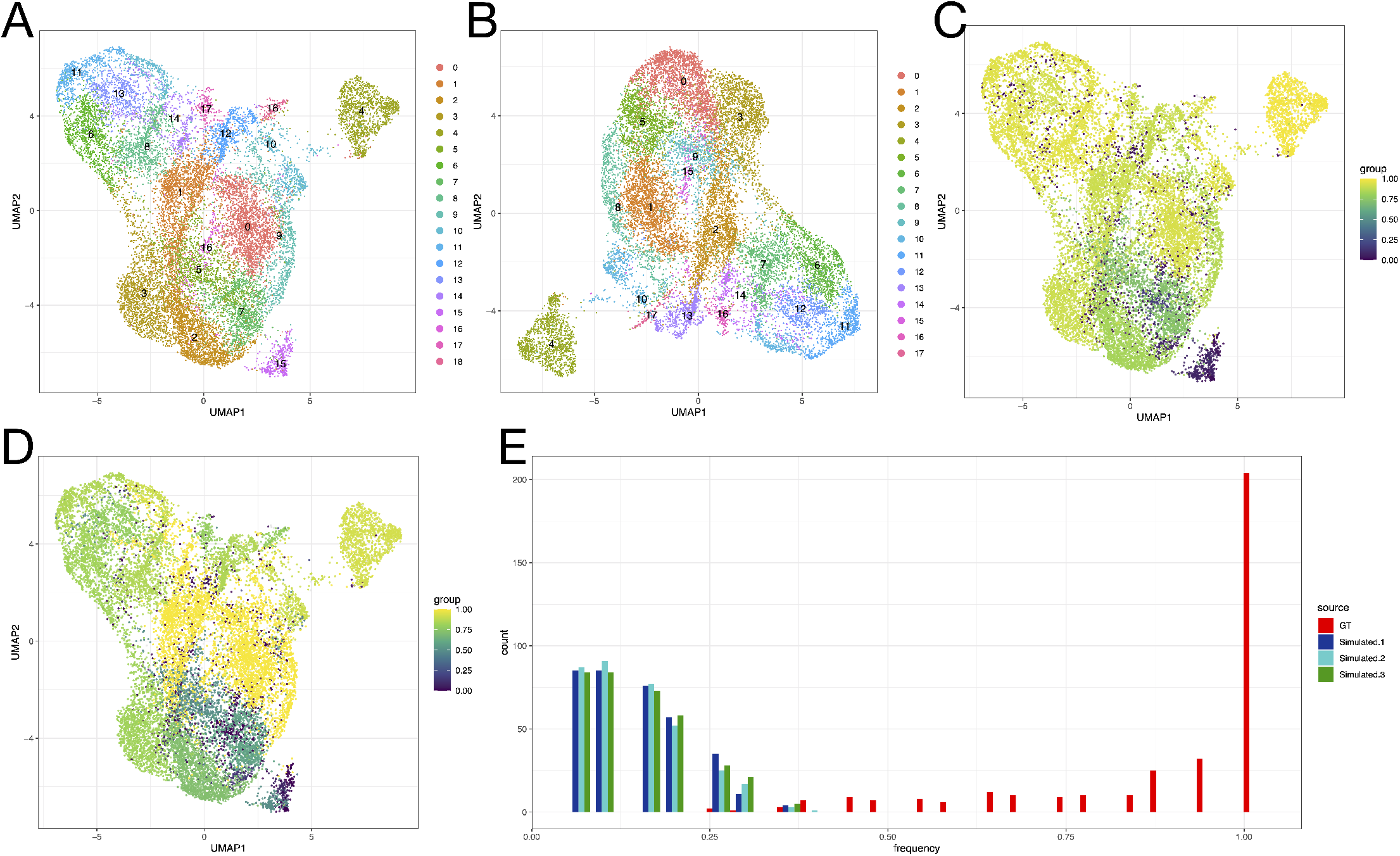
Lane splitting induces variability in clustering of P1 10x scRNA-seq data. A-B: UMAP representations of GT (A) and S (B) samples with colors indicating SLM clusters calculated on nearest-neighbor graphs; 19 clusters are found in GT and 18 in the S samples. C. Element-centric clustering similarity, highlighted using the colour gradient, reveals differences between clusterings at the bottom of the UMAP, especially for the lower right island of cells (vanishing island). D. Jaccard similarity of cluster markers across GT and S clusterings suggests differences in cell types inferred from the data. E. Fraction of within-island nearest neighbours for cells in the vanishing island in GT and 3 S samples. The island cells are no longer discrete from the larger body of cells in S samples.

To investigate whether the disappearing island was a reproducible effect of the lane splitting simulations, *n* = 3 S samples were generated. The S samples were consistent in terms of UMAP topography and SLM clustering results (Supplementary Fig 4A-F). Across all S samples the “island cells” were scattered in the larger body of cells (Supplementary Fig 4H). Conversely, in UMAPs generated on GT across 4 random seeds, the island of cells was consistently detached from the larger body of cells (Supplementary Fig 4G). Furthermore, the fraction of within-island nearest neighbors for the island cells was quantified in the GT and S samples (Fig 3E); while the island cells had each other as neighbors in GT, this is no longer observed in the simulations, suggesting the loss of some transcriptional heterogeneity during the lane splitting.

The marker genes that distinguish the vanishing island (Supplementary fig 5A) from the rest of the cells are SPP1, SRGN, SOCS2, ALDH1A1, AREG and HIST1H1C (Supplementary Fig 5B); in particular SPP1 appears highly specific to the island. Upon recalculating the PCA, and subsequently the UMAP, on the set of abundant genes with SPP1 excluded, the island is absorbed into the wider body of cells (Supplementary fig 5C), and so is SPP1 expression (Supplementary fig 5D). SPP1 is identified as the 5th most variable gene in GT by SCTransform. In S samples, marked differences in gene variance can be observed when compared to GT (Supplementary fig 5E), leading to the downstream consequences on dimensionality reductions and clustering results.

To further investigate the consequences of lane splitting clustering variability on cluster markers, typically used to infer the identity of cells, the per-cell Jaccard similarity index (JSI) was calculated (Fig 3D). Certain regions of cells, such as the middle right of the UMAP, displayed high JSI, indicating they would be interpreted similarly in the GT and S analyses. Other regions exhibited lower JSI; the vanishing island (cluster 15) had lower JSI than the retained island (cluster 4). These results suggest that the former could be interpreted inconsistently, depending on whether the library was split across lanes or not.

## Discussion

### Effects of splitting on read diversity and levels of noise

The main consequence of across-lane sample splitting is the variation in read diversity; this propagates onto transcript coverage and quantification. To assess the variation, we focused on the transcript complexity (defined as the ratio of unique to total reads [52], calculated per transcript), exemplified on the SRR7624365 sample (bulk mRNAseq, 0h rep 1). We compared the complexities, per transcript, of the GT and S samples (Fig 4A); each point corresponds to a gene, the colour gradient is proportional to the log_2_ (abn). We observe a consistent trend for medium-high abundant genes, across a wide complexity range [0, 0.75], highlighting the higher complexity (i.e. more diverse reads) of the GT sample. Also noticeable is the high variability in complexity for the low abundance genes, presented on the MA plots in the bulk section of the Results, and the localisation of the low abundance cloud under the equal complexity diagonal, enforcing the previous conclusion across all abundances.

**Figure 4:**
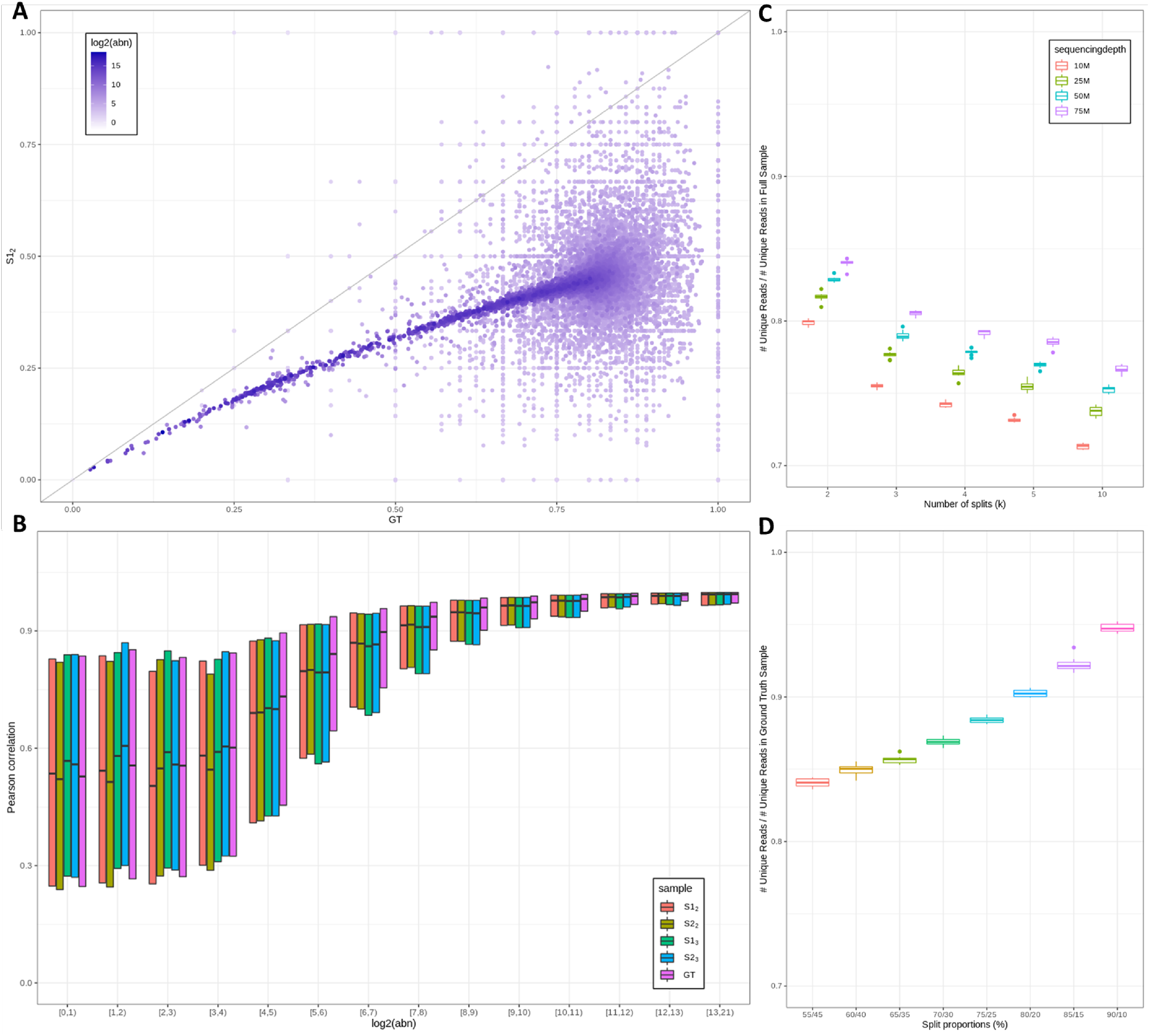
Summary of split-effects on the number of unique reads and noise. A) Scatter plot illustrating the complexity ratio (non-redundant to redundant counts) for the GT (x-axis) vs an S (y-axis) sample for SRR7624365 (bulk mRNAseq, 0h rep 1). Each point represents a gene and and the colour gradient is proportional to the log_2_ abundance. B) Box plot of the PCC binned by abundance for the transcript-based noise removal (noisyR applied to BAM files) corre-sponding to GT, *k* = 2 and *k* = 3 S samples for 0h rep 1. C) Boxplot showing distributions of ratios of recovered unique reads in the S samples with respect to the *k* hyper-parameter (*k* = 2, 3, 4, 5, 10); the effect of GT sample sequencing depth is also assessed. The distributions are built on 10 iterations. D) Boxplot showing distribution of ratios of recovered unique reads in the S samples when the proportions of concatenated subsamples are varied (from 55-45 to 90-10). The distributions are built on 10 iterations.

Yet another side effect of the variation in reads diversity is the quantification of noise across the samples [14]; we focused on the transcript-based approach in noisyR since it is directly influenced by the robustness of transcript coverage. In Fig 4B, we illustrate the variation of point-to-point PCC vs the variation in abundance for the GT and 2 iterations of *k* = 2 (S1_2_,S2_2_) and *k* = 3 (S1_3_,S2_3_). The wide and low PCC distributions correspond to higher levels of noise; the distributions become higher and tighter for medium to high abundance genes. For medium abundance genes, the GT distributions are systematically higher than *k* = 2 and *k* = 3, suggesting that the splitting increases the level of noise and thus interferes with the detection of DE genes or other downstream analyses, as illustrated in the Results section. The level of noise in the simulated samples is also variable and generally higher in *k* = 3 samples than *k* = 2, although the distribution of PCC is wider and lower for all S samples than GT.

### Effects of varying the number and proportions of splits

In the Results section, we exemplified the concatenation of *k* = 2 and *k* = 3 equal subsamples. However, in a real-world scenario, we rarely observe an equal number of reads across split samples; in addition, split-designs are occasionally mixed. To further understand the knock-on effects on downstream analyses, we illustrate the effect of variable number and proportions of reads allocated to splits. We used the bulk mRNAseq sample SRR7624365 (0h rep 1) as the starting point and, based on the consistency in conclusions across different inputs, we expect a convergence of results for other bulk or single-cell data. We subsampled the fastq files to 50%, 33%, 25%, 20% and 10% of the total sequencing depth and created S samples by concatenating the 2, 3, 4, 5 and 10 subsamples, respectively. For each case study, the distributions were generated on 10 iterations. We also assessed the sequencing depth co-variate; the results on the full sequencing depth (75.4M) and for samples subsampled to 50M, 25M and 10M reads respectively (without replacement) are presented. The ratios of recovered unique reads (i.e. number of unique reads in the S sample divided by the number of unique reads for the GT sample) across the simulated case studies are presented in Fig 4C. We observe a decrease in recovered ratios proportional to the number of splits and initial sequencing depth. The decrease in ratios is uniform across sequencing depths, but the recovered ratios also have lower y-intercepts (lower absolute ratios), highlighting that the effect of across lane-splitting is more extreme at low sequencing depths i.e. while the decrease-rate is lower for higher *k*-values, as number of splits (*k*) increases, we see a consistent reduction in the number of unique reads. This underlines that splitting across lanes has an adverse effect on the diversity of reads.

Additionally, we assessed the effect of uneven splits across lanes; focusing on *k* = 2 S samples, we varied the split proportions from 55-45 to 90-10. The resulting recovery ratios are shown in Fig 4D. The minimum for the recovery ratios is achieved for the 50-50 proportions; the recovery ratios gradually increases as the larger subsample approaches 100%. The observed increase is small for first few increments and increases more rapidly as proportions approach 100-0. This illustrates, from yet another angle, the variation in reads diversity.

On various sequencing datasets, bulk and single cell, we observed that the splitting of samples across lanes reduces the diversity of reads, which in turn triggers side effects on quantification (e.g. gene expression for mRNAseq, peak expression for ChIPseq) and auxiliary properties (such as the length of ChIP peaks). The splitting in itself introduces an additional level of variability in terms of robustness and reproducibility; it may pose added difficulties stemming from variable observed number of reads derived from the stochasticity of the sequencing itself. The potential batch effects derived from loading full samples on sequencing lanes can be mitigated through randomisation. We acknowledge that technical circumstances may make splitting unavoidable; our recommendation is a consistency in sequencing setup across all samples in an experiment.

## Funding

This research was supported by the Core grant awarded to the Wellcome-MRC Cambridge Stem Cells Institute (the Wellcome Trust [203151/Z/16/Z] and the UKRI Medical Research Council [MC PC 17230]).

## Acknowledgments

All authors acknowledge the constructive feedback from the Core Bioinformatics group. We thank Dr Chris Ellis for discussion on the splitting applied on 10x Genomics data. IM acknowledges early discussions of the idea that led to is project with Prof Tracey Chapman (School of Biological Sciences, University of East Anglia, NRP, UK) and Dr Rachel Rusholme-Pilcher (Earlham Institute, NRP, UK).

## Author contributions

EW performed the bulk analyses (mRNAseq and ChIPseq), including the subsampling design; RCG performed the smartSeq analyses; AS performed the 10x Genomics analyses. IM proposed and supervised the project. All authors wrote the manuscript, and read the final draft.

## Notes

### Competing Interest Statement

The authors have declared no competing interest.

### Summary of Updates

The effect of splitting samples across lanes was also illustrated on sequenced bulk mRNAseq data; this provided a data-driven justification for the subsampling approach (and a biological context for our study).

## References

[1] Stark R, Grzelak M, and Hadfield J. RNA sequencing: the teenage years. Nature Reviews Genetics, 20, 07 2019.

[2] Yandell M and Ence D. A beginner’s guide to eukaryotic genome annotation. Nature Reviews Genetics, 13:329–42, 04 2012.

[3] Steward C, Parker A, Minassian B, et al. Genome annotation for clinical genomic diagnostics: Strengths and weaknesses. Genome Medicine, 9, 05 2017.

[4] Salzberg S. Next-generation genome annotation: We still struggle to get it right. Genome Biology, 20, 12 2019.

[5] Conesa A, Madrigal P, Tarazona S, et al. A survey of best practices for RNA-seq data analysis. Genome Biology, 17, 01 2016.

[6] Oshlack A, Robinson M, and Young M. From RNA-seq reads to differential expression results. Genome Biology, 11:220, 12 2010.

[7] Lightbody G, Haberland V, Browne F, et al. Review of applications of high-throughput sequencing in personalized medicine: barriers and facilitators of future progress in research and clinical application. Briefings in Bioinformatics, 20(5):1795–1811, 06 2019.

[8] Lücken M and Theis F. Current best practices in single-cell RNA-seq analysis: a tutorial. Molecular Systems Biology, 15:e8746, 06 2019.

[9] McGuire A, Gabriel S, Tishkoff S, et al. The road ahead in genetics and genomics. Nature Reviews Genetics, 21:1–16, 08 2020.

[10] Stupnikov A, Tripathi S, de Matos Simoes R, et al. samExploreR: exploring reproducibility and robustness of RNA-seq results based on SAM files. Bioinformatics, 32(21):3345–3347, 07 2016.

[11] Schurch N, Schofield P, Gierliński M, et al. How many biological replicates are needed in an RNA-seq experiment and which differential expression tool should you use? RNA, 22, 03 2016.

[12] Oberg A, Bot B, Grill D, et al. Technical and biological variance structure in mrna-seq data: life in the real world. BMC Genomics, 13:304, 07 2012.

[13] Kim BS, Lee E, and Kim J. Analysis of technical and biological variability in single-cell RNA sequencing, volume 1935, pages 25–43. Humana Press, 01 2019.

[14] Moutsopoulos I, Maischak L, Lauzikaite E, et al. noisyR: enhancing biological signal in sequencing datasets by characterizing random technical noise. Nucleic Acids Research, 06 2021. gkab433.

[15] Ma X, Shao Y, Tian L, et al. Analysis of error profiles in deep next-generation sequencing data. Genome Biology, 20, 03 2019.

[16] Ross M, Russ C, Costello M, et al. Characterizing and measuring bias in sequence data. Genome Biology, 14:R51, 05 2013.

[17] K. Sorefan, Helio Pais, Adam Hall, Ana Kozomara, Sam Griffiths-Jones, Vincent Moulton, and Tamas Dalmay. Reducing ligation bias of small RNAs in libraries for next generation sequencing. Silence, 3, 05 2012.

[18] Hicks S, Townes FW, Teng M, and Irizarry R. Missing data and technical variability in single-cell RNA-sequencing experiments. Biostatistics (Oxford, England), 19, 11 2017.

[19] Reuter J, Spacek D, and Snyder M. High-throughput sequencing technologies. Molecular cell, 58:586–597, 05 2015.

[20] Eraslan G, Simon L, Mircea M, et al. Single-cell RNA-seq denoising using a deep count autoencoder. Nature Communications, 10:390, 01 2019.

[21] Chazarra-Gil R, Dongen S, Kiselev V, et al. Flexible comparison of batch correction methods for single-cell RNA-seq using BatchBench. Nucleic Acids Research, 49, 02 2021.

[22] Srivastava A, Malik L, Sarkar H, et al. Alignment and mapping methodology influence transcript abundance estimation. Genome Biology, 21:239, 09 2020.

[23] Dillies M, Rau A, Aubert J, et al. A comprehensive evaluation of normalization methods for Illumina high-throughput RNA sequencing data analysis. Briefings in Bioinformatics, 14, 09 2012.

[24] Mccarthy D, Chen Y, and Smyth G. Differential expression analysis of multifactor SRNA-Seq experiments with respect to biological variation. Nucleic Acids Research, 40:4288–97, 02 2012.

[25] Love M, Huber W, and Anders S. Moderated estimation of fold change and dispersion for RNA-Seq data with DESeq2. Genome Biology, 15:550, 12 2014.

[26] Svensson V, Natarajan K, Ly L, et al. Power analysis of single cell RNA-sequencing experiments. Nature Methods, 14, 03 2017.

[27] Nakato R and Shirahige K. Recent advances in ChIP-seq analysis: From quality management to whole-genome annotation. Briefings in Bioinformatics, 18:bbw023, 03 2016.

[28] Chung D, Kuan PF, Li B, et al. Discovering Transcription Factor Binding Sites in Highly Repetitive Regions of Genomes with Multi-Read Analysis of ChIP-Seq Data. PLoS Computational Biology, 7:e1002111, 07 2011.

[29] Dal Molin A and Camillo B. How to design a single-cell RNA-sequencing experiment: Pitfalls, challenges and perspectives. Briefings in Bioinformatics, 20, 01 2018.

[30] Mohorianu I, Bretman A, Smith D, et al. Genomic responses to socio-sexual environment in male drosophila melanogaster exposed to conspecific rivals. RNA, 23:rna.059246.116, 04 2017.

[31] Yang P, Humphrey SJ, Cinghu S, et al. Multi-omic profiling reveals dynamics of the phased progression of pluripotency. Cell Systems, 8(5):427–445.e10, 2019.

[32] Cuomo A, Seaton D, McCarthy D, et al. Single-cell RNA-sequencing of differentiating iPS cells reveals dynamic genetic effects on gene expression. Nature Communications, 11, 02 2020.

[33] Mende N, Bastos HP, Santoro A, et al. Quantitative and molecular differences distinguish adult human medullary and extramedullary haematopoietic stem and progenitor cell landscapes. bioRxiv, 2020.

[34] Thurmond J et al. FlyBase 2.0: the next generation. Nucleic Acids Research, 47(D1):D759–D765, 2018.

[35] Dobin A, Davis CA, Schlesinger F, et al. STAR: ultrafast universal RNA-seq aligner. Bioinformatics, 29(1):15–21, 10 2012.

[36] Liao Y, Smyth GK, and Shi W. featureCounts: an efficient general purpose program for assigning sequence reads to genomic features. Bioinformatics, 30(7):923–930, 11 2013.

[37] Ramírez F, Ryan DP, Grüning B, Bhardwaj V, Kilpert F, Richter AS, Heyne S, Dündar F, and Manke T. deepTools2: a next generation web server for deep-sequencing data analysis. Nucleic Acids Research, 44(W1):W160–W165, 04 2016.

[38] Yates AD, Achuthan P, Akanni W, et al. Ensembl 2020. Nucleic Acids Research, 48(D1):D682–D688, 11 2019.

[39] Bolstad BM, Irizarry RA, Åstrand M, et al. A comparison of normalization methods for high density oligonucleotide array data based on variance and bias. Bioinformatics, 19:185–193, 01 2003.

[40] Robinson M, Mccarthy D, and Smyth G. edgeR: a Bioconductor package for differential expression analysis of digital gene expression data. Bioinformatics, 26:139–140, 01 2010.

[41] Raudvere U, Kolberg L, Kuzmin I, et al. g:Profiler: a web server for functional enrichment analysis and conversions of gene lists (2019 update). Nucleic Acids Research, 47(W1):W191–W198, 05 2019.

[42] Zhang Y, Liu T, Meyer C, et al. Model-based analysis of ChIP-seq (MACS). Genome Biology, 9:R137, 10 2008.

[43] Andrews S, Krueger F, Segonds-Pichon A, et al. FastQC. Babraham Institute, January 2012.

[44] Ewels P, Magnusson M, Lundin S, et al. MultiQC: summarize analysis results for multiple tools and samples in a single report. Bioinformatics, 32(19):3047–3048, 06 2016.

[45] Stuart T, Butler A, Hoffman P, et al. Comprehensive integration of single cell data. Cell, 11 2018.

[46] Hafemeister C and Satija R. Normalization and variance stabilization of single-cell RNA-seq data using regularized negative binomial regression. Genome Biology, 20, 12 2019.

[47] Beckers M, Mohorianu I, Stocks M, et al. Comprehensive processing of high throughput small RNA sequencing data including quality checking, normalization and differential expression analysis using the UEA sRNA Workbench. RNA, 23:rna.059360.116, 03 2017.

[48] Zheng G, Terry J, Belgrader P, et al. Massively parallel digital transcriptional profiling of single cells. Nature Communications, 8:14049, 01 2017.

[49] Waltman L and van Eck NJ. A smart local moving algorithm for large-scale modularity-based community detection. European Physical Journal B, 86, 08 2013.

[50] Gates A, Wood I, Hetrick W, et al. Element-centric clustering comparison unifies overlaps and hierarchy. Scientific Reports, 9:8574, 06 2019.

[51] Shahsavari A and Mohorianu I. ClustAssess: Tools for Assessing Clustering, 2021. R package version 0.1.1.

[52] Mohorianu I, Schwach F, Jing R, et al. Profiling of short RNAs during fleshy fruit development reveals stage-specific sRNAome expression patterns. The Plant journal : for cell and molecular biology, 67:232–46, 03 2011.

