## Supplementary figures for "The sum of two halves may be different from the whole. Effects of splitting sequencing samples across lanes"

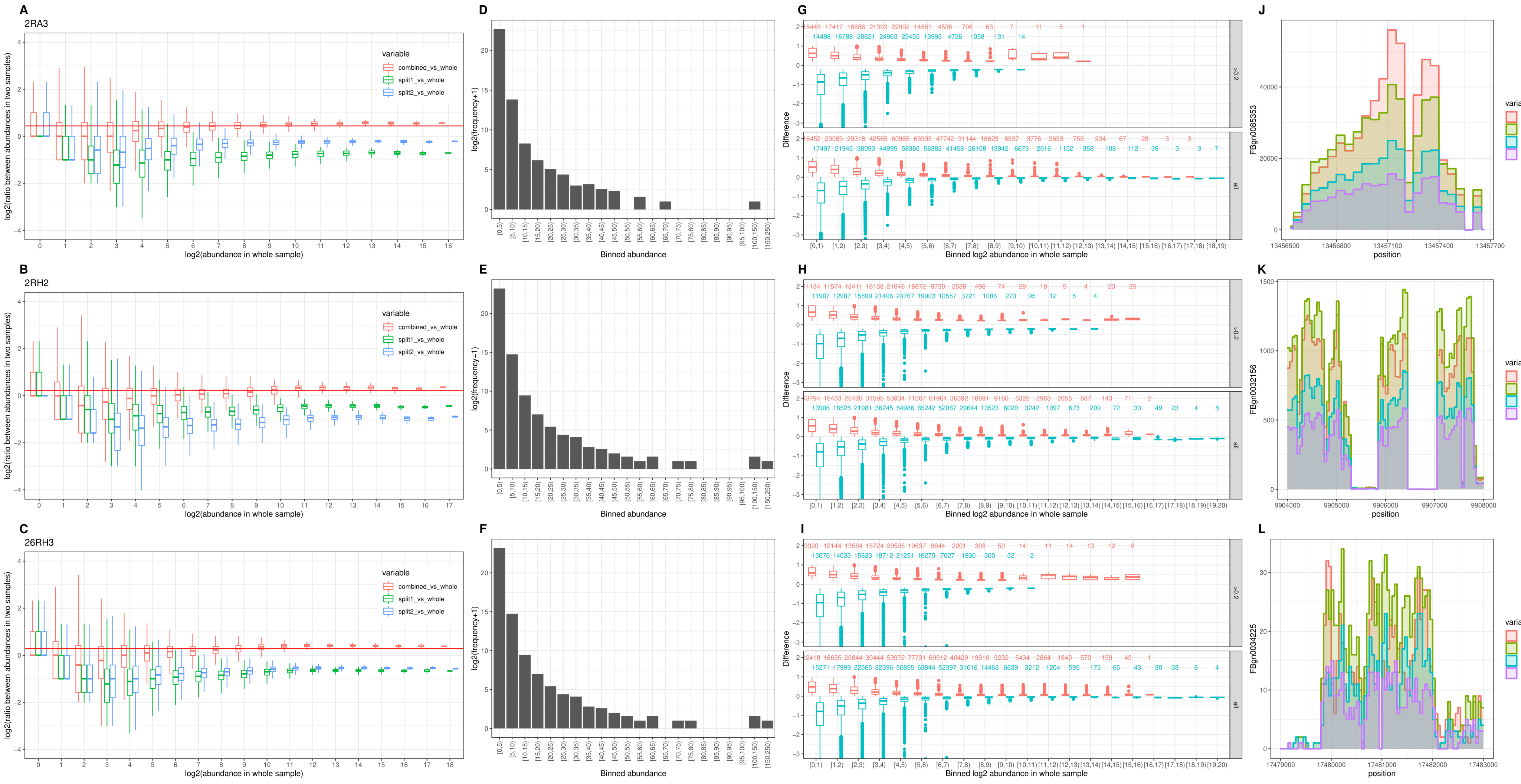


Supplementary figure 1: Differences observed in . *melanogaster* bulk mRNAseq samples sequenced on either one lane or split across two lanes

*Supplementary figure 1:* ***Assessment of differences for bulk mRNAseq samples sequenced on either one lane or split across two lanes***

*A-C. Boxplot showing log ratio of redundant counts in separate (split 1 and split 2) and combined split sample (concatenated) and whole sample across log2(abundances) for the fragments which are non-zero in the split and whole samples. The red line shows the ratio between the sequencing depth of the combined split sample and the whole sample and is where the ratios are expected to fall. We see underrepresentation of low-medium abundances and overamplification of high abundances in the split sample compared to the whole.*

*D-F. Barplot showing distribution of redundant count for fragments found only in the whole sample, and so lost in the split sample, capped at abundance 250. While the majority of lost fragments have low abundance, there are some medium-high abundances also lost. G-I. Positive and negative differences between average abundance in whole and combined split sample across windows of length 100 over transcripts. The abundance in the combined split is scaled by the ratio between sequencing depths and differences are scaled by average abundance across the window in the whole sample. All windows (bottom) and only windows with scaled difference of ¿0.2 (top) are considered. A positive (+) difference means the average abundance over the window was higher in the whole sample and negative (-) the converse. The number of windows are shown above each box. The first row (A,D,G) corresponds to sample 2RA3, the second (B,E,H) to 2RH2 and the third (C,F,I) to 26RH3. J-L. The examples of transcripts from 2RH2 with (J) high abundance (FBgn0085353), (K) medium abundance (FBgn0032156) and (L) low abundances (FBgn0034225), showing the distribution of abundance along the transcript in the whole, split and combined split samples.*

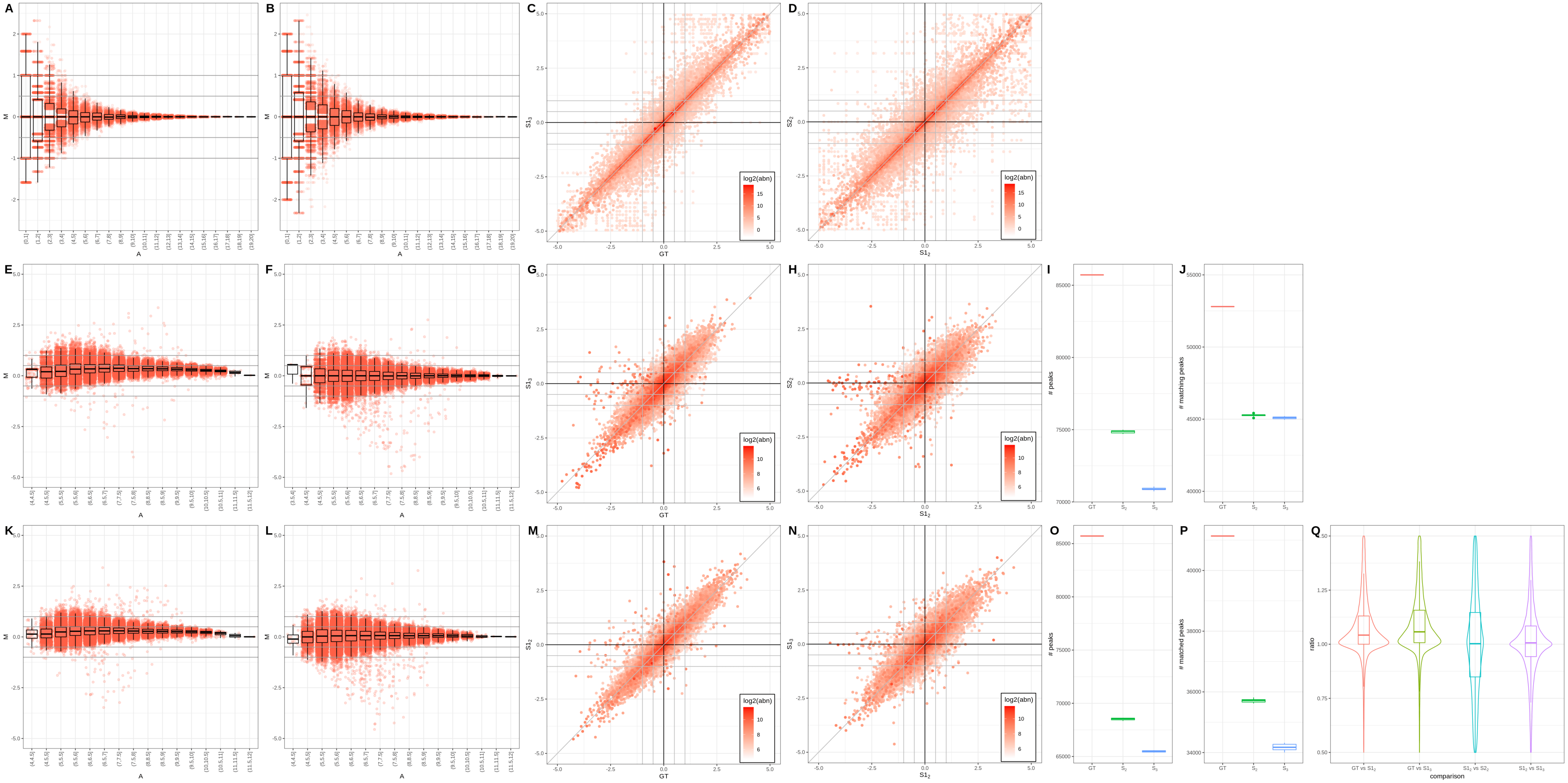

Supplementary figure 2: Bulk mRNA-seq results illustrating the quantification stability across GT and S samples.

*Supplementary figure 2:*

***Bulk mRNA-seq results illustrating the quantification stability across GT and S samples.***

*A-B. Scatter (MA) and box plot showing $log_2$ abundance against $log_2FC$ within A) GT sample (SRR7624365, 0h rep 1) and a $k = 3$ S sample and B) two $k = 2$ S samples for the same sample (SRR7624365, 0h rep 1). In both cases, we see the standard MA funneling shape, however a shift above to y-axis is observed, resulting from the lack of signal for low abundance genes for GT vs $k = 3$ (A). The distribution for $k = 2$ vs $k = 2$ is more symmetrical but wider, illustrating the technical variation introduced by splitting.*

*C-D. Scatter (cross) plot showing $log_2(FC)$ when comparing C) 0h vs 12h DE for GT samples and k=2 S samples, and D) 0h vs 12h DE for 2 k=2 S samples; the colour gradient is proportional to the average abundance across samples. Overall the DE predictions converge (genes are close to the diagonal); a larger variability in FC is observed for $k = 2$ vs $k = 2$ than GT vs $k = 3$.*

***Bulk H3K4me3 ChIPseq results illustrating the quantification stability across GT and S samples.***

*E-F. Scatter (MA) and box plot showing $log_2$ average abundance against $|log_2FC|$ within E) GT sample (SRR7624381, 0h) and a $k = 3$ sample $S1_3$ and F) two $k = 2$ S samples ($S1_2$,$S2_2$) for the same sample (SRR7624381, 0h). We see a shift above the x-axis for low amplitude peaks (borderline vanishing peaks) for GT vs $k = 3$ (E) and an over-amplification of peaks, from the wide variability at medium amplitude.*

*G-H. Cross plot on $log_2FC$s for G) 0h vs 12h DE for GT samples and k=2 S samples, and H) 0h vs 12h DE for 2 $k = 2$ S samples; the colour gradient is proportional to the average abundance of genes. We see several "false negative" peaks in the simulated samples (appearing along the central horizontal range) due to borderline vanishing peaks.*

*I) Boxplot showing number of peaks matched between 0h and 12h samples, in GT sample, for $n = 10$ iterations of $k = 2$ ($S1_2$) and $k = 3$ ($S1_3$) samples. The number of peaks is noticeably lower in the S samples compared to GT; the observed effect is stronger for $k = 3$ than $k = 2$ i.e. peaks are lost through sample splitting.*

*J) Boxplot showing number of peaks in GT sample (SRR7624381, 0h), for $n = 10$ iterations of $k = 2$ ($S1_2$) and $k = 3$ ($S1_3$) samples. As well as a reduction in the number of peaks called per sample, we also see a reduction in the number of peaks which can be matched between time-points and thus can be used for differential methylation calls.*

***Comparison of H3K27ac ChIPseq analysis results for GT and S samples.***

*K-L. Scatter (MA) and box plots showing $log_2$ abundance against $log_2$ FC for K) GT sample (SRR7624389, 0h) and a $k = 2$ S sample and L) $k = 2$ and $k = 3$ S samples. We see a shift towards positive M values for low amplitude peaks (borderline vanishing peaks) for GT vs $k = 3$ (K) and an over-amplification of peaks, from the wide variability at medium amplitude.*

*M-N. Scatter (cross) plot on $|log_2FC|$s when comparing M) 0h vs 12h DE for GT samples and $k = 2$ S samples, and N) 0h vs 12h DE for $k = 2$ and $k = 3$ S samples; the colour gradient is proportional to the average amplitude across samples. We see several false negative peaks in simulated data (appearing along the central horizontal channel) due to borderline vanishing peaks.*

*O) Boxplot showing number of peaks matched between 0h and 12h samples, in GT sample, $n = 10$ iterations of $k = 2$ ($S_2$) and $k = 3$ ($S_3$) samples. The number of peaks is significantly smaller in the simulated samples compared to GT, and the peak count is smaller for $k = 3$ than $k = 2$. This underlines that peaks are lost by splitting samples.*

*P) Boxplot showing number of peaks in GT sample (SRR7624389, 0h), $n = 10$ iterations of $k = 2$ ($S_2$) and $k = 3$ ($S_3$) samples. As well as a reduction in the number of peaks called per sample, we also see a reduction in the number of peaks which can be matched between time-points and thus used for differential methylation calls.*

*Q) Violin and box plots showing distribution of ratio of peak lengths between 2 samples (sample 1 peak length / sample 2 peak length), comparing GT $k = 2$ vs S sample ($S1_2$ and $S2_2$) and $k = 3$ S samples ($S1_3$). We notice a global, systematic shift towards shorter peaks in simulated samples; the effect is more pronounced for $k = 3$. The between-simulation comparison distributions are symmetrical but the $k = 2$ vs $k = 2$ distribution is unexpectedly wide.*

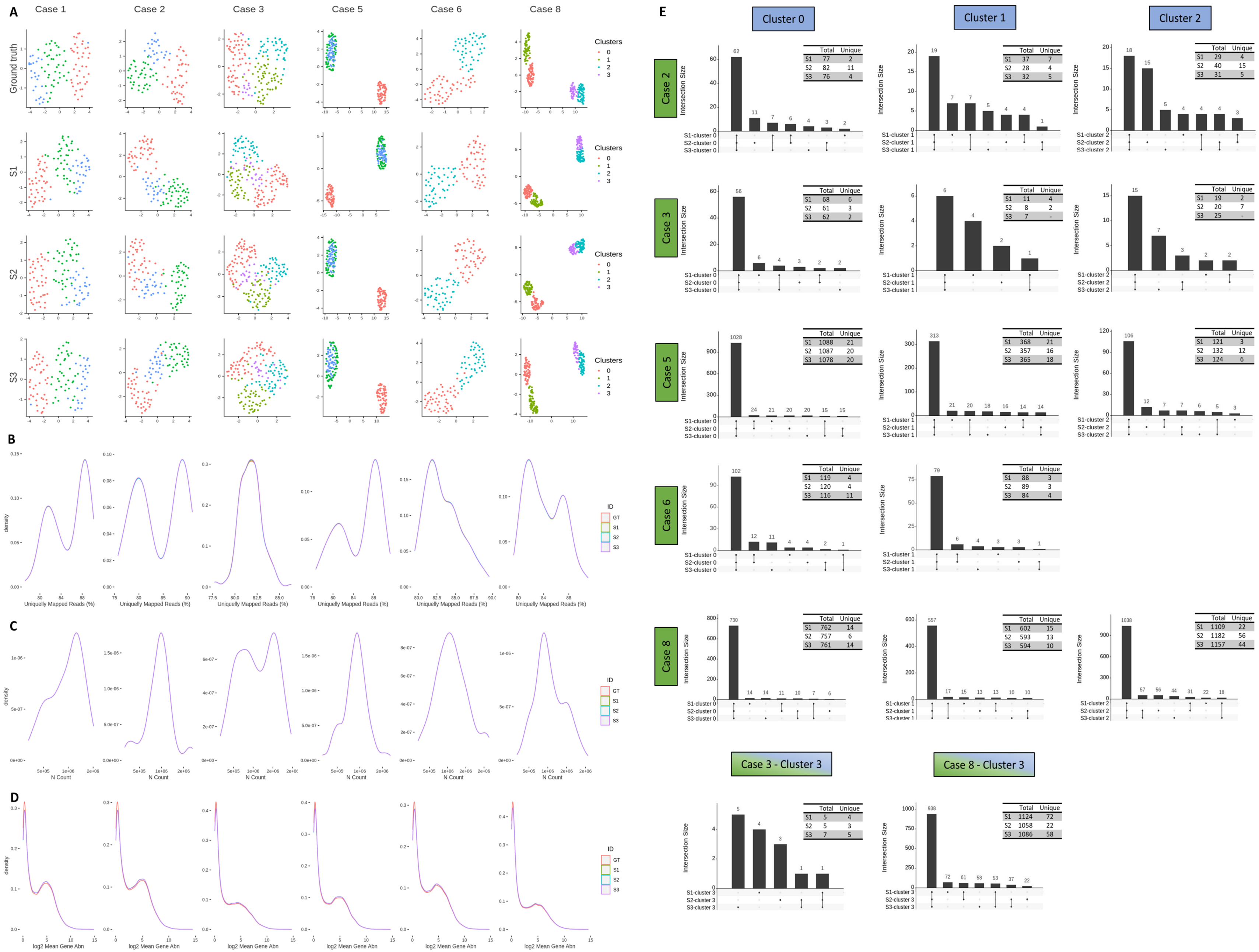


Supplementary figure 3: Lane splitting influences cell topology and feature abundances without affecting the fraction of uniquely mapped reads or the count distribution. In addition, it introduces variability across replicates.

*Supplementary figure 3:* ***Lane splitting influences cell topology and feature abundances without affecting the fraction of uniquely mapped reads or the count distribution. In addition, it introduces variability across replicates.***

*For the different study cases considered (Table 1) we show: A) UMAP representation of the GT and S samples. Cells are coloured by their cluster identity. Sample splitting results in modifications of the data topology that are visible in low dimensional representations of the data. B) Fraction of Uniquely Mapped Reads distribution, illustrating the consistency in signal for the GT and S samples. C) Number of counts distribution for the GT and S samples, illustrating the split approach i.e. the number of reads in the GT and S samples match. D) Mean gene abundance, across the GT and S samples, distribution in $log_2$ scale. E) Consistency in DE analysis on the smartSeq data. Sequencing lane splitting introduces a degree of variability across replicates that propagates to downstream analyses e.g. DE. The upset plots summarise the intersection of the DE features across clusters for the S samples, for each study case considered. The adjacent table to each plot shows the total and unique number of DE entries detected for each set of S samples.*

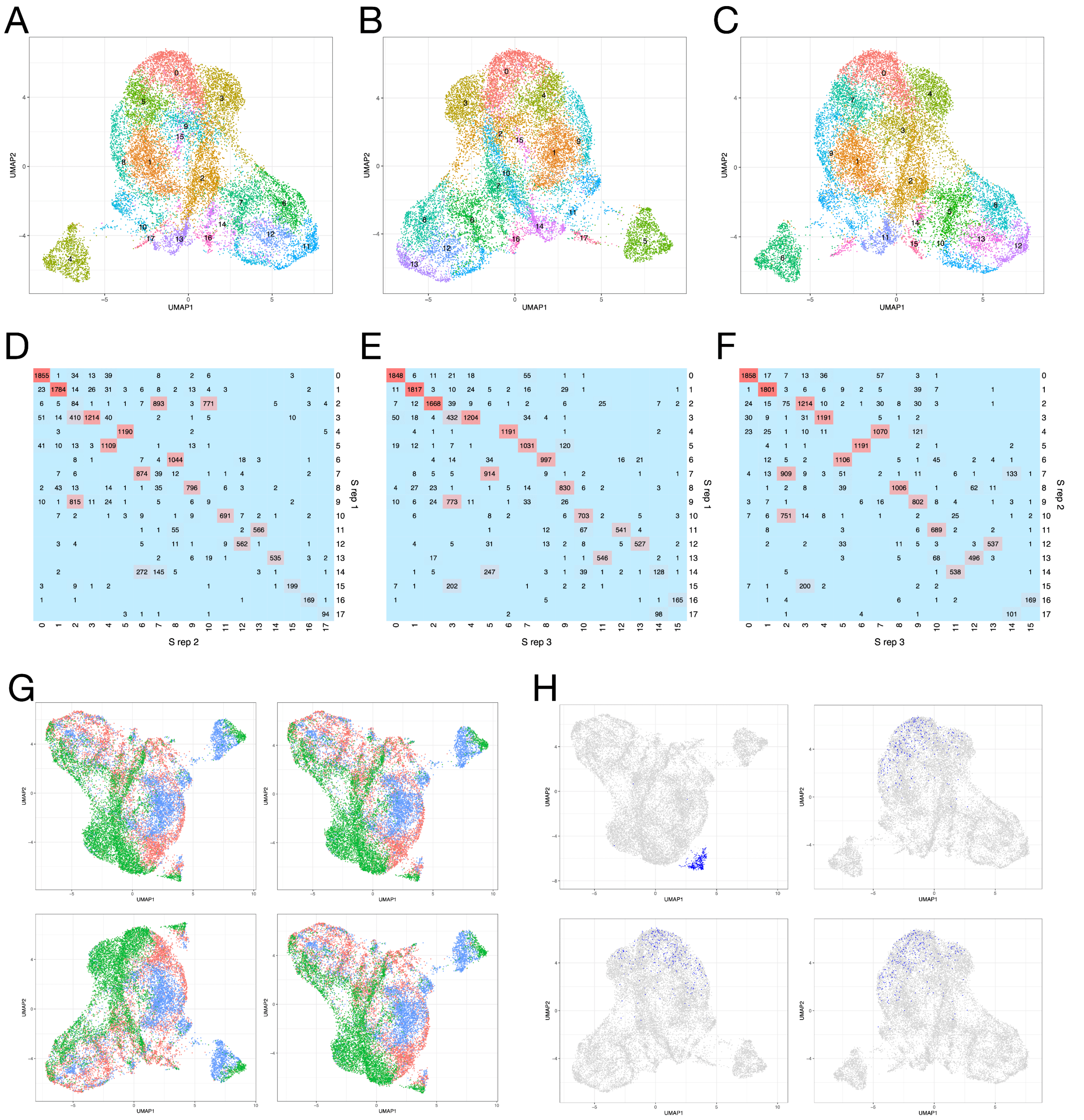


Supplementary figure 4: **Consistency between replicate 10x S samples clustering; vanishing island of cells in GT is assimilated in S samples.**

UMAPs of 3 S samples, generated using consecutive random seeds, illustrate consistency between simulations (A-C); colors indicate SLM clusters. UMAPs display similar general topography across random seeds, with rotation being the main difference. High agreement is also observed in pairwise contingency tables of cell assignments for clusterings across random seeds (D for clusterings A vs B; E for A vs C; F for B vs C), with almost one-to-one relationship between clusters in a majority of comparisons. G. UMAPs of GT samples, generated across 4 consecutive random seeds, illustrate a group of cells (upper right of top left subpanel) consistently separating from the larger body of cells. Colors indicate tissue origin of cells (red: bone marrow, green: peripheral blood, blue: spleen).

H. UMAPs of GT (top left subpanel) and 3 S samples, with blue indicating cells belonging to cluster 15 (vanishing island) from the GT samples. The cluster of cells is distinct in GT, but absorbed by the larger body of cells in each of the simulations.

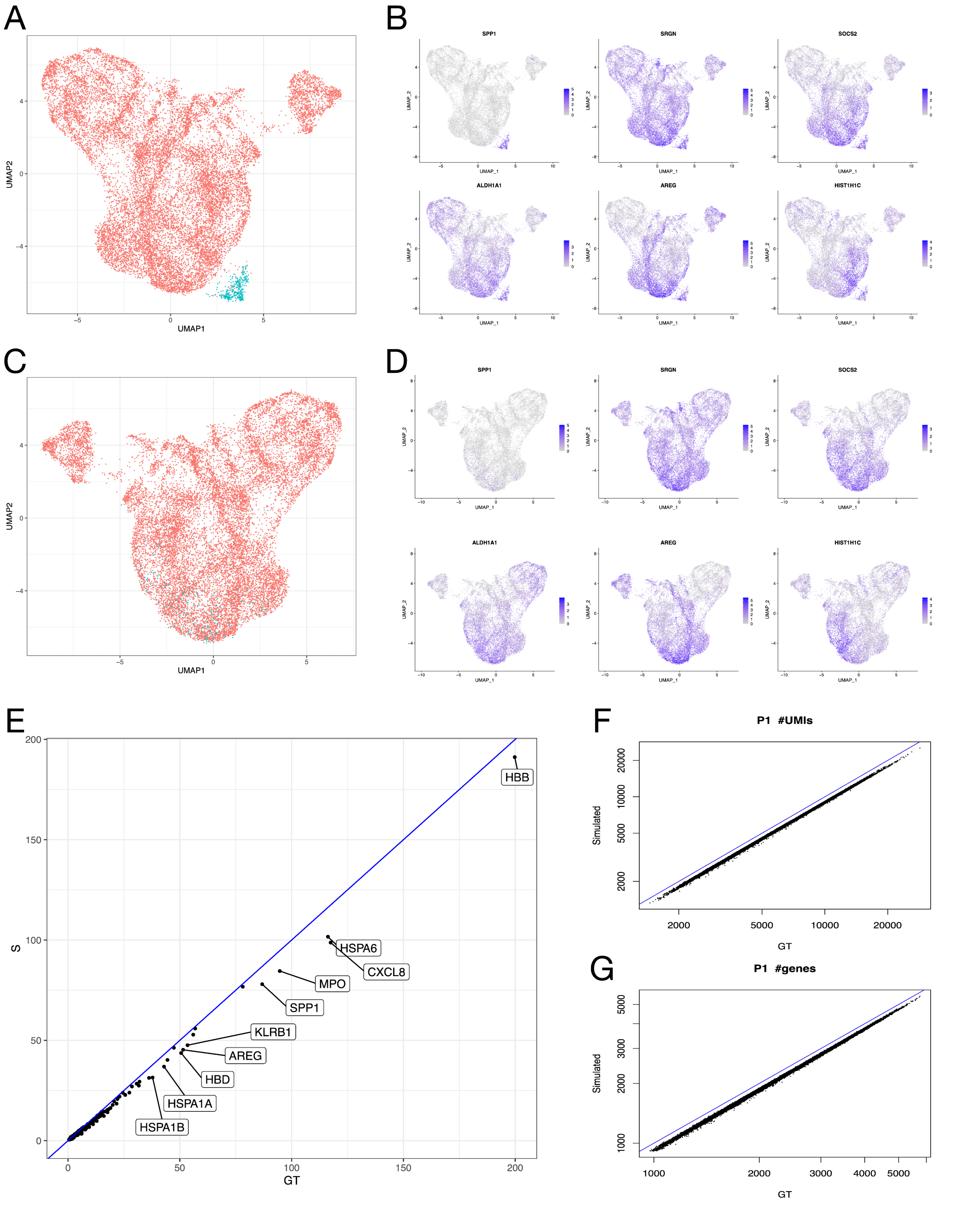


Supplementary figure 5: The vanishing island depends strongly on the SPP1 expression; the simulations induce transcriptome-wide changes in UMI diversity and gene diversity and variability.

*Supplementary figure 5:* ***The vanishing island depends strongly on the SPP1 expression; the simulations induce transcriptome-wide changes in UMI diversity and gene diversity and variability.***
*A-B. The vanishing island cells (A) have 6 markers genes with* $|FC| > 2$ *(B) The expression UMAPs for these markers illustrate the localisation of signal; the color gradient is proportional to the logarithm of the abundance. SPP1 is particularly specific for the island.*
*C-D. The PCA and UMAP were recalculated on the set of abundant genes excluding SPP1; the island cells were absorbed into the wider body of cells (C). SPP1 expression is now more widely spread at the bottom of the UMAP (D). We infer that the vanishing island depends strongly on SPP1 expression.*
*E. The most variable genes have lower variance in S samples (the ten genes with greatest absolute difference in variance between GT and S are labeled). In addition, the set of the 3,000 most variable genes, used by default for dimensionality reductions and clustering by SCTransform, is different between GT and S (intersect of 2,704 genes). The geometric mean of variance ratios illustrates slight transcriptome-wide decrease of gene variance in S (GT increase vs S of 0.63%). Blue line corresponds to 1:1 ratio.*
*F-G. Scatter plots of the number of UMIs (F) and number of unique genes per cell (G) reveal lower transcriptomic resolution in S samples. Each point corresponds to one cell. GT samples consistently have more UMIs and higher number of unique genes per cell. The geometric mean of ratios indicates that lane splitting consistently leads to loss of transcriptomic information (GT increase vs S of 11.6% for UMIs, 7.7% for genes). Blue line corresponds to 1:1 ratio. Note the axes are on log-scale.*

### Table legends

***Supplementary table 1*** *Tables of* $\chi^2$ *p-values comparing the per-position nucleotide content in the whole, split and combined split samples for [A] 2RA3, [B] 2RH2, [C] 26RH3 with Benjamini-Hochberg multiple testing correction.*
***Supplementary table 2*** *[A] P-values of the Kolmogorov-Smirnoff test on distributions of the: Number of Unique Reads, Fraction of Uniquely Mapped Reads, Number of Counts, Number of Features and mean Gene Abundances; for the GT vs S comparisons. [B] Kullback-Leibler Divergence values on distributions of the: Number of Unique Reads, Fraction of Uniquely Mapped Reads, Number of Counts, Number of Features and mean Gene Abundances; for the GT vs S comparisons.*
